# Next generation sequencing to investigate genomic diversity in Caryophyllales

**DOI:** 10.1101/646133

**Authors:** Boas Pucker, Tao Feng, Samuel F. Brockington

**Author notes:** corresponding author: Boas Pucker.

## Abstract

Caryophyllales are a highly diverse and large order of plants with a global distribution. While some species are important crops like *Beta vulgaris*, many others can survive under extreme conditions. This order is well known for the complex pigment evolution, because the pigments anthocyanins and betalains occur with mutual exclusion in species of the Caryophyllales. Here we report about genome assemblies of *Kewa caespitosa* (Kewaceae), *Macarthuria australis* (Macarthuriaceae), and *Pharnaceum exiguum* (Molluginaceae) which are representing different taxonomic groups in the Caryophyllales. The availability of these assemblies enhances molecular investigation of these species e.g. with respect to certain genes of interest.

## Introduction

Caryophyllales form one of the largest flowering plant order and are recognized for their outstanding ability to colonise extreme environments. Examples are the evolution of Cactaceae in deserts, extremely fast radiation [1–3] e.g. in arid-adapted Aizoaceae and in carnivorous species in nitrogen-poor conditions. Caryophyllales harbor the greatest concentration of halophytic plant species and display repeated shifts to alpine and arctic habitats in Caryophyllaceae and Montiaceae. Due to these extreme environments, species exhibit many adaptations [2–4] such as specialized betalain pigments to protect photosystems in high salt and high light conditions [5]. There are several examples for repeated evolution in the Caryophyllales e.g. leaf and stem succulence for water storage, various mechanisms for salt tolerance, arid-adapted C_4_ and CAM photosynthesis [4], and insect trapping mechanisms to acquire nitrogen [6].

In addition, to their fascinating trait evolution, the Caryophyllales are well known for important crops and horticultural species like sugar beet, quinoa and spinach. Most prominent is the genome sequence of *Beta vulgaris* [7] which was often used as a reference for studies within Caryophyllales [7–10]. In addition, genomes of *Spinacia oleracea* [7,11], *Dianthus caryophyllus* [12], *Amaranthus hypochondriacus* [13], and *Chenopodium quinoa* [14] were sequenced. Besides *Carnegiea gigantea* and several other cacti [15], recent genome sequencing projects were focused on crops due to their economical relevance. However, genome sequences of other species within the Caryophyllales, are needed to provide insights into unusual patterns of trait evolution.

The evolution of pigmentation is known to be complex within the Caryophyllales [8] with a single origin of betalain and at least three reversals to anthocyanin pigmentation. The biosynthetic pathways for betalain and anthocyanin pigmentation are both well characterized. While previous studies have demonstrated that the genes essential for anthocyanin synthesis persists in betalain pigmented taxa [16,17], the fate of the betalain pathway in the multiple reversals to anthocyanin pigmentations is unknown. Here, we sequenced three species from different families to contribute to the genomic knowledge about Caryophyllales: *Kewa caespitosa* (Kewaceae), *Macarthuria australis* (Macarthuriaceae), and *Pharnaceum exiguum* (Molluginaceae) were selected as representatives of anthocyanic lineages within the predominantly betalain pigmented Caryophyllales. *K. caespitosa* and *P. exiguum* are examples of putative reversals from betalain pigmentation to anthocyanic pigmentation, while *Macarthuria* is a lineage that diverged before the inferred origin of betalain pigmentation [8].

Several transcript sequences of the three plants investigated here were assembled as part of the 1KP project [18]. Since the sampling for this transcriptome project was restricted to leaf tissue, available sequences are limited to genes expressed there. Here we report three draft genome sequences to complement the available gene set and to enable analysis of untranscribed sequences like promoters, regulatory elements, pseudogenes, and transposable elements.

## Material & Methods

### Plant material

The seeds of *Kewa caespitosa* (Friedrich) Christenh., *Marcarthuria australis* Hügel ex Endl., and *Pharnaceum exiguum* Adamson were obtained from Millennium Seed Bank (London, UK) and were germinated at the Cambridge University Botanic Garden. The plants were grown in controlled glasshouse under conditions: long-day (16 h light and 8 h dark), 20 °C, 60% humidity. About 100 mg fresh young shoots were collected and immediately frozen in liquid nitrogen. Tissue was ground in liquid nitrogen using a mortar and pestle. DNA was extracted using the QIAGEN DNeasy Plant Mini Kit (Hilden, Germany) and RNA was removed by the QIAGEN DNase-Free RNase Set. DNA quantity and quality were assessed by Nanodrop (Thermofisher Scientific, Waltham, MA, USA) and agarose gel electrophoresis. DNA samples were sent to BGI Technology (Hongkong) for library construction and lllumina sequencing.

### Sequencing

Libraries of *K. caespitosa, M. australis*, and *P. exiguum* were sequenced on an Illumina HiSeq X-Ten generating 2×150nt reads (AdditionalFile 1). Trimmomatic v0.36 [19] was applied for adapter removal and quality trimming as described previously [20]. Due to remaining adapter sequences, the last 10 bases of each read were clipped. FastQC [21] was applied to check the quality of the reads.

### Genome size estimation

The size of all three investigated genomes was estimated based on k-mer frequencies in the sequencing reads. Jellyfish v2 [22] was applied for the construction of a k-mer table with parameters described by [23]. The derived histogram was further analyzed by GenomeScope [23] to predict a genome size. This process was repeated for all odd k-mer sizes between 17 and 25 (AdditionalFile 2). Finally, an average value was selected from all successful analyses.

### Genome assembly

The performance of different assemblers on the data sets was tested (AdditionalFile 3, AdditionalFile 4, AdditionalFile 5). While CLC Genomics Workbench performed best for the *M. australis* assembly, SOAPdenovo2 [24] showed the best results for *K. caespitosa* and *P. exiguum* and was therefore selected for the final assemblies. To optimize the assemblies, different k-mer sizes were tested as this parameter can best be adjusted empirically [25]. First, k-mer sizes from 67 to 127 in steps of 10 were evaluated, while most parameters remained on default values (AdditionalFile 6). Second, assemblies with k-mer sizes around the best value of the first round were tested. In addition, different insert sizes were evaluated without substantial effect on the assembly quality. In accordance with good practice, assembled sequences shorter than 500 bp were discarded prior to downstream analyses. Custom Python scripts [20,26] were deployed for assembly evaluation based on simple statistics (e.g. N50, N90, assembly size, number of contigs), number of genes predicted by AUGUSTUS v3.2 [27] *ab initio*, average size of predicted genes, and number of complete BUSCOs [28]. Scripts are available on github: https://github.com/bpucker/GenomeAssemblies2018.

BWA-MEM v0.7 [29] was used with the –m flag to map all sequencing reads back against the assembly. REAPR v1.0.18 [30] was applied on the selected assemblies to identify putative assembly errors through inspection of paired-end mappings and to break sequences at those points.

The resulting assemblies were further polished by removal of non-plant sequences. First, all assembled sequences were subjected to a BLASTn [31] against the sugar beet reference genome sequence RefBeet v1.5 [7,32] and the genome sequences of *Chenopodium quinoa* [14], Carnegiea gigantea [15], *Amaranthus hypochondriacus* [13], and *Dianthus caryophyllus* [12]. Hits below the e-value threshold of 10^−10^ were considered to be of plant origin. Second, all sequences without hits in this first round were subjected to a BLASTn search against the non-redundant nucleotide database nt. Sequences with strong hits against bacterial and fungal sequences were removed as previously described [20,26]. BLASTn against the *B. vulgaris* plastome (KR230391.1, [33]) and chondrome (BA000009.3, [34]) sequences was performed to identify and remove sequences from these organelle subgenomes.

### Assembly quality assessment

Mapping of sequencing reads against the assembly and processing with REAPR [30] was the first quality control step. RNA-Seq reads (AdditionalFile 7) were mapped against the assemblies to assess completeness of the gene space and to validate the assembly with an independent data set. STAR v2.5.1b [35] was used for the RNA-Seq read mapping as previously described [26].

### Genome annotation

RepeatMasker [36] was applied using crossmatch [37] to identify and mask repetitive regions prior to gene prediction. Masking was performed in sensitive mode (-s) without screening for bacterial IS elements (-no_is) and skipping interspersed repeats (-noint). Repeat sequences of the Caryophyllales (-species caryophyllales) were used and the GC content was calculated per sequence (-gccalc). Protein coding sequences of transcriptome assemblies (AdditionalFile 7) were mapped to the respective genome assembly via BLAT [38] to generate hints for the gene prediction process as previously described [39]. BUSCO v3 [28] was deployed to optimize species-specific parameter sets for all three species based on the sugar beet parameter set [40]. AUGUSTUS v.3.2.2 [27] was applied to incorporate all available hints with previously described parameter settings to optimize the prediction of non-canonical splice sites [39]. Different combinations of hints and parameters were evaluated to achieve an optimal annotation of all three assemblies. A customized Python script was deployed to remove all genes with premature termination codons in their CDS or spanning positions with ambiguous bases. Representative transcripts and peptides per locus were identified based on maximization of the encoded peptide length. INFERNAL (cmscan) [41] was used for the prediction of non-coding RNAs based on models from Rfam13 [42].

Functional annotation was transferred from *Arabidopsis thaliana* (Araport11) [43] via reciprocal best BLAST hits as previously described [26]. In addition, GO terms were assigned to protein coding genes through an InterProScan5 [44]-based pipeline [26].

### Comparison between transcriptome and genome assembly

The assembled genome sequences were compared against previously published transcriptome assemblies (AdditionalFile 7) in a reciprocal way to assess completeness and differences. BLAT [38] was used to align protein coding sequences against each other. This comparison was limited to the protein coding sequences to avoid biases due to UTR sequences, which are in general less reliably predicted or assembled, respectively [39]. The initial alignments were filtered via filterPSL.pl [45] based on recommended criteria for gene prediction hint generation to remove spurious hits and to reduce the set to the best hit per locus e.g. caused by multiple splice variants.

## Results

### Genome size estimation and genome sequence assembly

Prior to the *de novo* genome assembly, the genome sizes of *Kewa caespitosa, Macarthuria australis*, and *Pharnaceum exiguum* were estimated from the sequencing reads (Table 1, AdditionalFile 1). The estimated genome sizes range from 265 Mbp (*P. exiguum*) to 623 Mbp (*M. caespitosa*). Based on these genome sizes, the sequencing coverage ranges from 111x (*K. caespitosa*) to 251x (*M. australis*).

**Table 1:** Genome size estimation and *de novo* assembly statistics.

|  | *Kewa caespitosa* | *Macarthuria australis* | *Pharnaceum exiguum* |
| --- | --- | --- | --- |
| Accession | GCA_900322205 | GCA_900322265 | GCA_900322385 |
| Estimated genome size [Mbp] | 623 | 497.5 | 265 |
| Sequencing coverage | 111x | 251x | 206x |
| Assembly size (-N) | 531,205,354 | 525,292,167 | 254,526,612 |
| Number of sequence | 55,159 | 271,872 | 16,641 |
| N50 | 28,527 | 2,804 | 56,812 |
| Max. sequence length | 340,297 | 211,626 | 514,701 |
| GC content | 38.1% | 36.6% | 37.4% |
| Complete BUSCOs | 83.6% | 44.4% | 84.3% |
| Assembler | SOAPdenovo2 | CLC Genomics<br>Workbench v9 | SOAPdenovo2 |
| k-mer size | 79 | Automatic | 117 |

Different assembly tools and parameters were evaluated to optimize the assembly process (AdditionalFile 3, AdditionalFile 4, AdditionalFile 5). Sizes of the final assemblies ranged from 254.5 Mbp (*P. exiguum*) to 531 Mbp (*K. caespitosa*) (Table 1, AdditionalFile 8). The best continuity was achieved for *P. exiguum* with an N50 of 57 kbp.

### Assembly validation

The mapping of sequencing reads against the assembled sequences resulted in mating rates of 99.5% (*K. caespitosa*), 98% (*M. australis*), and 94.8% (*P. exiguum*). REAPR identified between 1390 (*P. exiguum*) and 16181 (*M. australis*) FCD errors which were corrected by breaking assembled sequences. The mapping of RNA-Seq reads to the polished assembly resulted in mapping rates of 53.9% (*K. caespitosa*) and 43.1% (*M. australis*), respectively, when only considering uniquely mapped reads. Quality assessment via BUSCO revealed 83.6% (*K. caespitosa*), 44.4% (*M. australis*), and 84.3% (*P. exiguum*) complete benchmarking universal single copy ortholog genes (n=1440). In addition, 6.5% (*K. caespitosa*), 21.7% (*M. australis*), and 4.0% (*P. exiguum*) fragmented BUSCOs as well as 9.9% (*K. caespitosa*), 33.9% (*M. australis*), and 11.7% (*P. exiguum*) missing BUSCOs were identified. The proportion of duplicated BUSCOs ranges from 1.5% (*K. caespitosa*) to 2.1% (*P. exiguum*). The number of duplicated BUSCOs was high in *M. australis* (11.8%) compared to both other genome assemblies (1.5% and 2.1%, respectively).

### Genome annotation

After intensive optimization (AdditionalFile 9), the polished structural annotation contains between 26,155 (*P. exiguum*) and 80,236 (*M. australis*) protein encoding genes per genome (Table 2). The average number of exons per genes ranged from 2.9 (*M. australis*) to 6.6 (*K. caespitosa*). Predicted peptide sequence lengths vary between 241 (*M. australis*) and 447 (*K. caespitosa*) amino acids. High numbers of recovered BUSCO genes support the assembly quality (Fig. 1). Functional annotations were assigned to between 50% (*K. caespitosa*) and 70% (*P. exiguum*) of the predicted genes per species. These assemblies revealed between 598 (P. exiguum) and 1604 (M. australis) putative rRNA, 821 (*K. caespitosa*) to 1492 (*M. australis*) tRNA genes, and additional non-protein-coding RNA genes (Table 2).

**Fig. 1.**
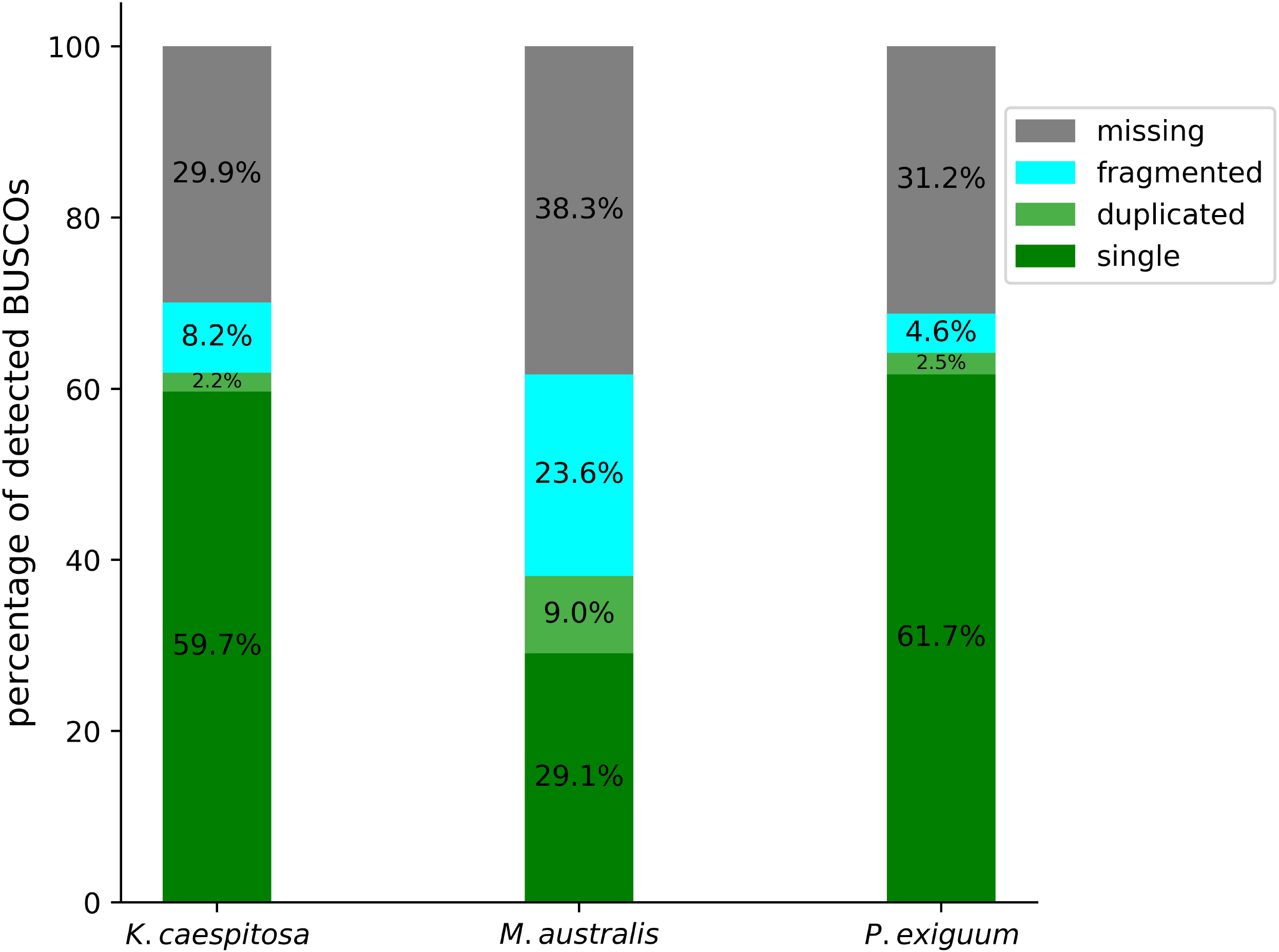
Assembly completeness. Assembly completeness was assessed based on the proportion of complete, fragmented, and missing BUSCOs.

**Table 2:** Assembly annotation statistics. These data sets are available at https://doi.org/10.4119/unibi/2941041 (Kewa caespitosa), https://doi.org/10.4119/unibi/2941043 (Macarthuria australis), and https://doi.org/10.4119/unibi/2941045 (Pharnaceum exiguum).

|  | *Kewa caespitosa* | *Macarthuria australis* | *Pharnaceum exiguum* |
| --- | --- | --- | --- |
| Final gene number | 50661 | 80236 | 26,155 |
| Functional annotation assigned | 25,058 (49.46%) | 50,536 (62.98%) | 18,372 (70.24%) |
| Average gene lengths [bp] | 5494 | 1936 | 5090 |
| Average mRNA length [bp] | 2143 | 1018 | 2154 |
| Average peptide length [aa] | 447 | 241 | 435 |
| RBHs vs. BeetSet2 | 9,968 | 10,568 | 10,045 |
| Average number of exons per gene | 6.6 | 2.9 | 6 |
| Number of predicted tRNAs | 821 | 1491 | 1260 |
| Number of predicted rRNAs | 720 | 1604 | 598 |

### Comparison between transcriptome and genome assemblies

Previously released transcriptome assemblies were compared to the genome assemblies to assess completeness and to identify differences. In total 44,169 of 65,062 (67.9%) coding sequences of *the K. caespitose* transcriptome assembly were recovered in the corresponding genome assembly. This recovery rate is lower for both *M. australis* assemblies, where only 27,894 of 58,953 (47.3%) coding sequences were detected in the genome assembly. The highest rate was observed for *P. exiguum*, where 37,318 of 42,850 (87.1%) coding sequences were found in the genome assembly. When screening the transcriptome assemblies for transcript sequences predicted based on the genome sequences, the recovery rate was lower (Fig. 2). The number of predicted representative coding sequences with best hits against the transcriptome assembly ranged from 16.3% in *K. caespitosa* to 29.7% in *P. exiguum* thus leaving most predicted coding sequences without a good full length hit in the transcriptome assemblies.

**Fig. 2.**
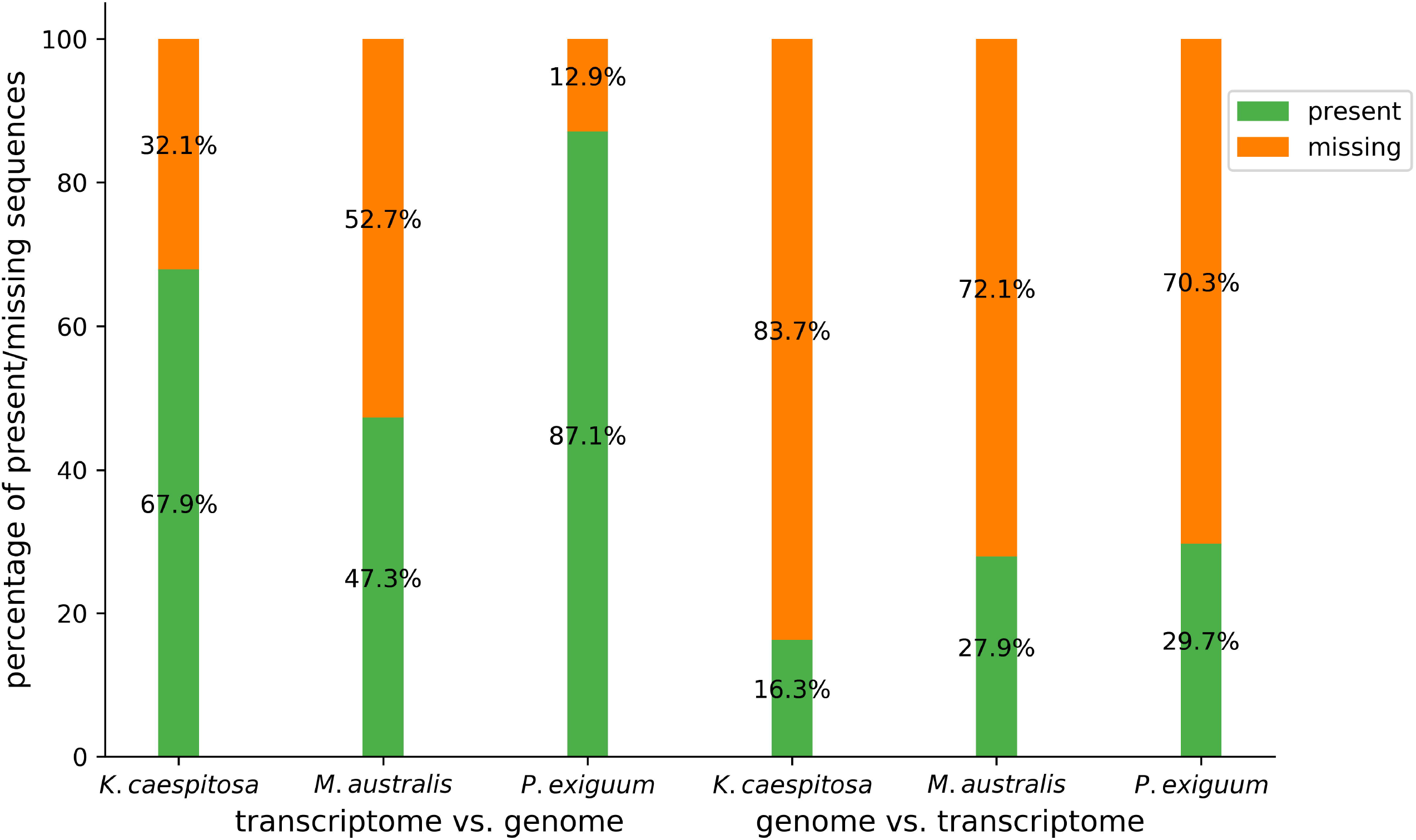
Recovery of sequences between transcriptome and genome assemblies. The figure displays the percentage of sequences present in one assembly that are recovered or missing in the other assembly type.

## Discussion

An almost perfect match between the predicted genome size and the final assembly size was observed for *P. exiguum*. When taking gaps within scaffolds into account the *K. caespitosa* assembly size reached the estimated genome size. High heterozygosity could be one explanation for the assembly size exceeding the estimated haploid genome size of *M. australis*. The two independent genome size estimations for *M. australis* based on different read data sets indicate almost perfect reproducibility of this method. Although centromeric regions and other low complexity regions were probably underestimated in the genome size estimation as well as in the assembly process, this agreement between estimated genome size and final assembly size indicates a high assembly quality. The continuity of the *P. exiguum* assembly is comparable to the assembly continuity of *Dianthus caryophyllus* [12] with a scaffold N50 of 60.7 kb. Additional quality indicators are the high proportion of detected BUSCOs in the final assemblies as well as the high mapping rate of reads against the assemblies. The percentage of complete BUSCOs is in the same range as the value of the *D. caryophyllales* genome assembly which revealed 88.9% complete BUSCOs based on our BUSCO settings. We demonstrate a cost-effective generation of draft genome assemblies of three different plant species. Investing into more paired-end sequencing based on Illumina technology would not substantially increase the continuity of the presented assemblies. This was revealed by initial assemblies for *M. australis* performed with less than half of all generated sequencing reads. Although the total assembly size increased when doubling the amount of incorporated sequencing reads, the continuity is still relatively low. No direct correlation between the sequencing depth and the assembly quality was observed in this study. Genome properties seem to be more influential than the amount of sequencing data. Even very deep sequencing with short reads in previous studies [12,20] was unable to compete with the potential of long reads in genome assembly projects [13,14], No major breakthroughs were achieved in the development of publicly available short read assemblers during the last years partly due to the availability of long reads which made it less interesting.

The number of predicted genes in *P. exiguum* is in the range expected for most plants [46,47], While the predicted gene numbers for *K. caespitosa* and *M. australis* are much higher than that for *P. exiguum*, they are only slightly exceeding the number of genes predicted for other plants [46,47], Nevertheless, the assembly continuity and the heterozygosity of *M. australis* are probably the most important factors for the artificially high number of predicted genes. The high percentage of duplicated BUSCOs (11.8%) indicates the presence of both alleles for several genes. As the average gene length in *M. australis* is shorter than in both other assemblies, some gene model predictions might be too short. This gene prediction could be improved by an increase in assembly continuity.

There is a substantial difference between the transcriptome sequences and the predicted transcripts of the genome assembly. The presence of alternative transcripts and fragmented transcripts in the transcriptome assemblies are one explanation why not all transcripts were assigned to a genomic locus. Another explanation is the intraspecific variation, as the genome and transcriptome assemblies were generated using different individuals. Some transcripts probably represent genes which are not properly resolved in the genome assemblies. This is especially the case for *M. australis*. The high percentage of complete BUSCOs of the *K. caespitosa* and *P. exiguum* genome assemblies indicate that missing sequences in the genome assemblies account only for a minority of the differences. The complete BUSCO percentage of the *P. exiguum* genome assembly even exceeds the value assigned to the corresponding transcriptome assembly. Although BUSCOs are selected in a robust way, it is likely that some of these genes are not present in the genomes investigated here, since *B. vulgaris* is the closest relative with an almost completely sequenced genome [7]. Our genome assemblies provide additional sequences of genes which are not expressed in the tissues sampled for the generation of the transcriptome assembly. In addition, coding sequences might be complete in the genome assemblies, while low expression caused a fragmented assembly based on RNA-Seq reads. This explains why only a small fraction of the predicted coding sequences of the genome assemblies was mapped to the coding sequences derived from the corresponding transcriptome assembly.

The availability of assembled sequences as well as large sequencing read data sets enables the investigation of repeats e.g. transposable elements across a large phylogenetic distance within the Caryophyllales. It also allows the extension of genome-wide analysis, such as gene family investigations from *B. vulgaris* to more representatives across Caryophyllales. As all three species produce anthocyanins, we provide the basis to study the underlying biosynthetic genes. Due to the huge evolutionary distance to other anthocyanin producing species, the availability of these sequences could facilitate the identification of common and unique features of the involved enzymes.

## Supporting information

AdditionalFile 1

AdditionalFile 2

AdditionalFile 3

AdditionalFile 4

AdditionalFile 5

AdditionalFile 6

AdditionalFile 7

AdditionalFile 8

AdditionalFile 9

## Author contribution

TF isolated DNA. BP and TF performed data processing, assembly, and annotation. BP, TF, and SFB interpreted the results. BP wrote the initial draft. All authors read and approved the final version of this manuscript.

## Acknowledgements

We thank the CeBiTec Bioinformatic Resource Facility team for great technical support.

## Supporting Information

**AdditionalFile 1. Sequencing result overview.**

**AdditionalFile 2. Genome size estimation results.** Genome size estimations with GenomeScope [23] are listed for various k-mer sizes. Two different read sets of *M. australis* were used for the genome size estimation (1=ERR2401802, 2=ERR2401614) to check the reproducibility.

**AdditionalFile 3. Evaluation of assembly attempts of *K. caespitosa*.**

**AdditionalFile 4. Evaluation of assembly attempts of *M. australis*.**

**AdditionalFile 5. Evaluation of assembly attempts of *P. exiguum*.**

**AdditionalFile 6. Detailed list of assembly parameters.**

**AdditionalFile 7. Gene prediction hint sources.** These RNA-Seq read data sets and transcriptome assemblies were incorporated in the gene annotation process as hints.

**AdditionalFile 8. Assembly attempt evaluation results.** Statistics of raw assemblies were calculated to identify the best parameter settings. Since k-mer size was previously reported as the most important parameter, extensive optimization was performed. In addition, different settings for insert sizes were evaluated for *P. exiguum* (phe001-phe006). Parameter optimization for *M. australis* was performed on a subset of all reads due to availability.

**AdditionalFile 9. Gene prediction statistics.** Different gene prediction approaches were performed during the optimization process. Results of these predictions include *ab initio* gene prediction and hint-based approaches. RNA-Seq reads and coding sequences derived from previous transcriptome assemblies are two incorporated hint types. In addition, we assessed the impact of repeat masking prior to gene prediction.

