## Supplementary figures and images for "Next generation sequencing to investigate genomic diversity in Caryophyllales"

### AdditionalFile 3

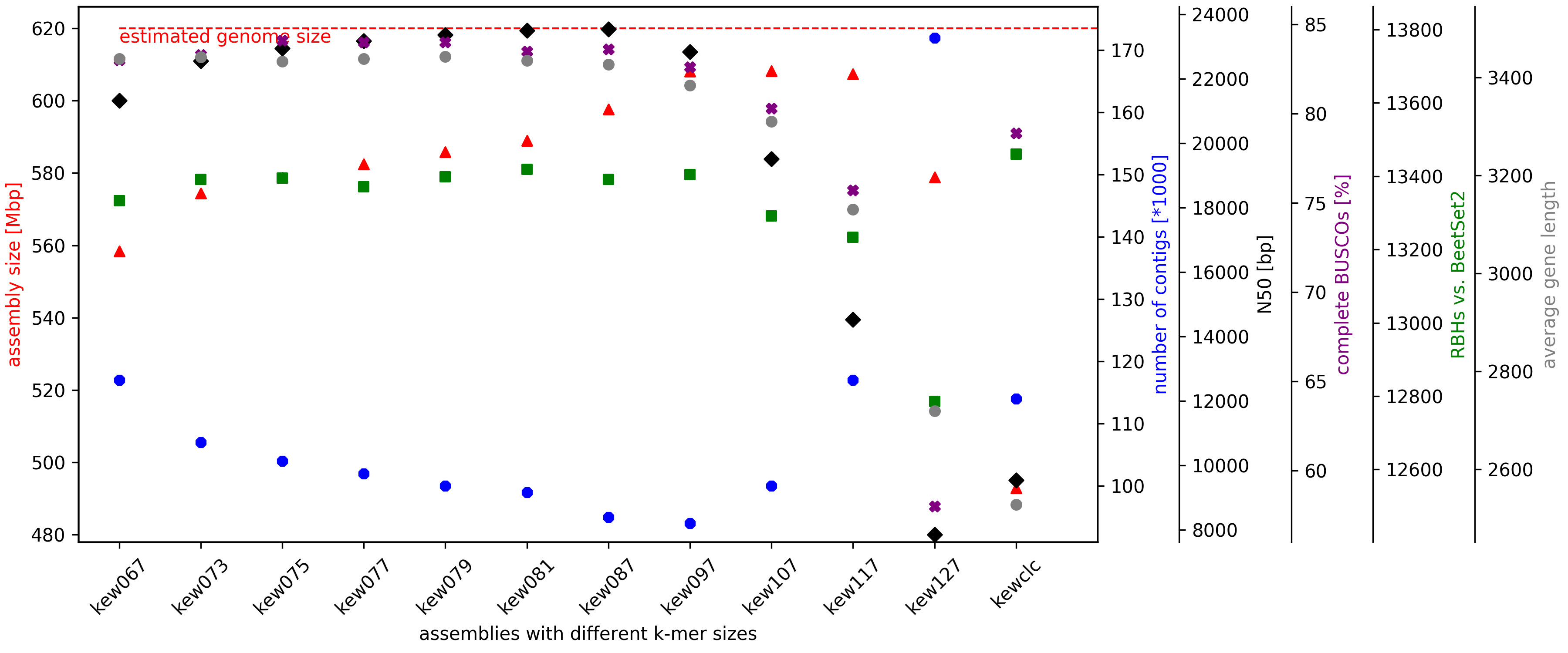

### AdditionalFile 4

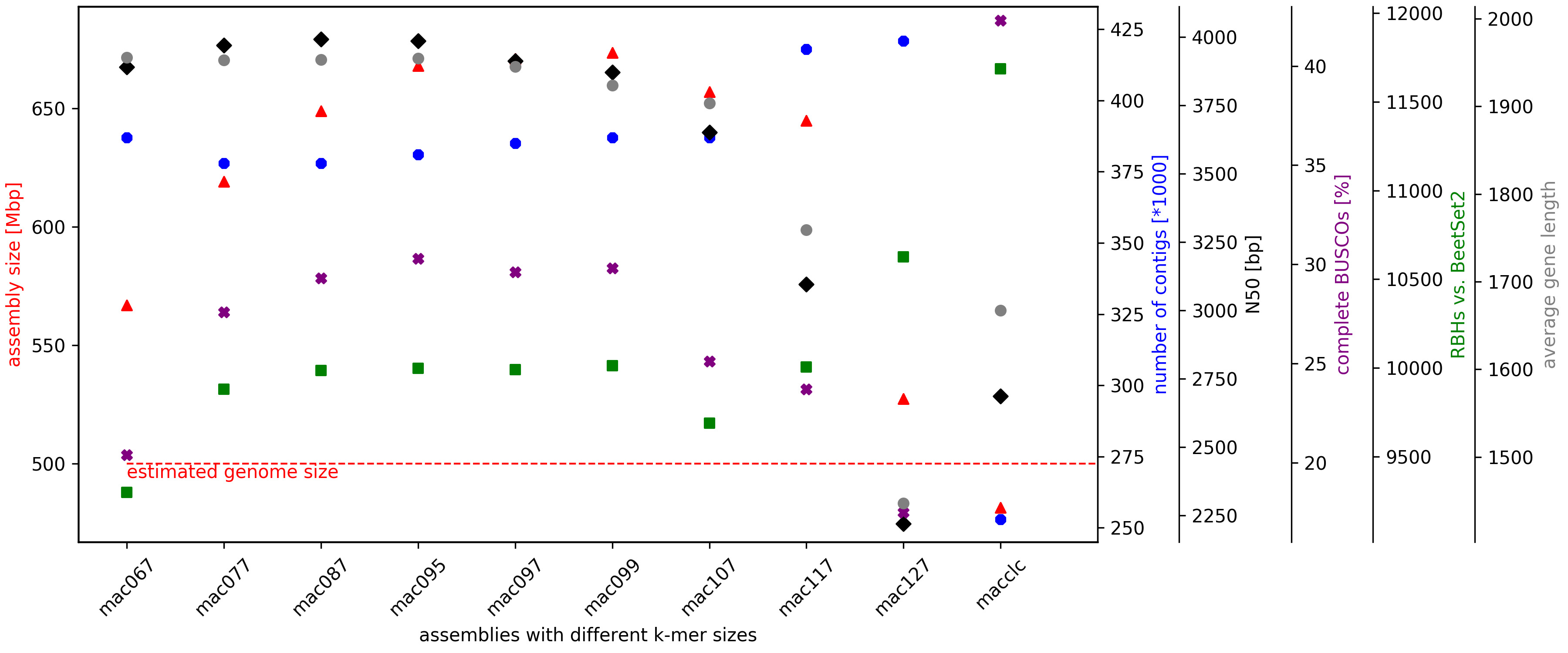

### AdditionalFile 5

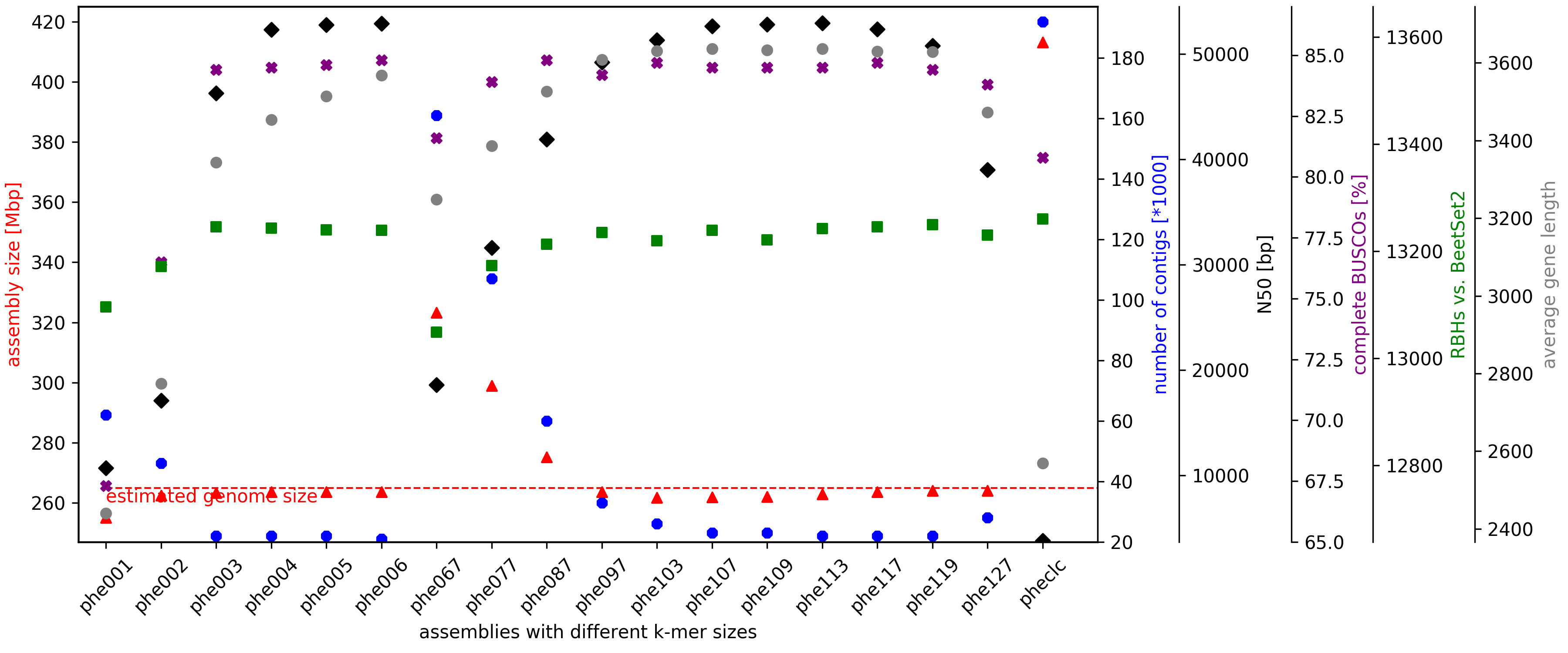
